# Investigation of In Vivo Silk Scaffold Degradation by Decoupling Tissue Ingrowth Using a GPR-Driven Digital Twin Framework

**DOI:** 10.64898/2026.06.10.731278

**Authors:** Guikang Wang, Yanjiu Li, Zhuowei Shen, Xianying Chen, Shiwen Zheng, Yaqin Li, Jianliu Wang, Xiuli Sun, Di Jia

## Abstract

Pelvic organ prolapse (POP) reconstruction is increasingly performed utilizing knitted silk meshes (KSM), yet tracking in vivo degradation kinetics remains challenging due to complex host tissue integration. This study developed an AI-driven semi-empirical framework utilizing Gaussian Process Regression (GPR) to bridge the kinetic mismatch between in vitro and in vivo environments. KSM scaffolds underwent 32 weeks of accelerated in vitro enzymatic degradation, with morphology (SEM), molecular conformation (FTIR), and mass loss being coupled with mechanical decay to train the GPR model. In vitro results revealed a multi-stage physical disintegration via a topochemical erosion pathway that preserved crystalline β-sheet structures despite macro-scale mass and mechanical loss. When validated in a rat abdominal wall defect model, traditional tracking metrics encountered severe bottlenecks. Heterogeneous dye labeling caused premature fluorescence quenching by Week 16, while extensive tissue ingrowth masked gravimetric and SEM signatures. Intriguingly, a bi-phasic in vivo mechanical trajectory was identified, where initial degradation-led failure was followed by a secondary mechanical recovery driven by biomechanical synergy with neo-muscular tissue. Importantly, despite premature quenching, this work presents the first optical imaging approach to visually mapping the complete chronological breakdown of the scaffold’s peripheral boundary layer in vivo, proving that outer functionalized layers eroded prior to internal silk cores. Furthermore, our GPR framework elegantly resolved the perennial technical barrier of tissue–mesh overlapping. By mathematically decoupling intrinsic polymer degradation from confounding tissue ingrowth, the model successfully achieved a first-of-its-kind prediction of the bare scaffold’s long-term structural fate in a non-adhered state, providing a robust digital twin methodology for lifetime predictions of degradable biomaterials.

## Introduction

Pelvic organ prolapse (POP) remains a critical global health crisis that severely compromises the psychological well-being, anatomical integrity, and overall quality of life for millions of women worldwide [1-3]. For decades, synthetic polymeric meshes, most notably polypropylene (PP), have served as the clinical gold standard for pelvic floor reconstruction. However, transvaginal PP implants frequently provoke severe, irreversible long-term complications, including severe chronic inflammatory responses, host tissue erosion, and catastrophic mechanical mismatch arising from the rigid, non-compliant nature of synthetic plastics[4, 5]. To transcend these clinical limitations, knitted silk meshes (KSM) derived from Bombyx mori have gained substantial traction as a highly competitive biopolymer alternative. This structural interest is primarily driven by silk fibroin’s exceptional biological compatibility and its unique crystalline β-sheet domain hierarchy, which imparts a remarkable mechanical resilience that perfectly mimics the anisotropic elastomeric profiles of native fascial tissues[6-12]. For successful functional tissue restoration, the structural degradation kinetics of the prosthetic matrix must be meticulously synchronized with the physiological rate of host neo-tissue deposition to guarantee a seamless, complication-free transition of load-bearing responsibilities[13, 14]. However, the hyper-dynamic and enzymatically hostile microenvironment of the pelvic cavity renders the real-time in vivo structural fate of KSM virtually “invisible” and highly unpredictable through conventional animal experimentation or standard clinical observation modalities.

This tracking bottleneck is fundamentally multi-faceted and currently introduces critical experimental blind spots into longitudinal biomaterial evaluations. On one hand, while non-invasive near-infrared (NIR) fluorescence tracking is widely celebrated as an optical standard for in vivo profiling[15], it frequently encounters premature signal extinction artifacts that misrepresent the actual material lifespan[16]. This premature extinction stems from a distinct topological phenomenon: due to the high steric hindrance and dense packing architecture of pristine fibroin bundles, the exogenous fluorophores predominantly functionalize only the superficial boundary fiber layers. This peripheral localization triggers a dense spatial self-quenching effect (De-quenching effect) that rapidly fades as the outer shell erodes, completely failing to reflect the continuous degradation of the unlabeled interior crystalline core matrix[17]. On the other hand, traditional destructive characterizations, such as gravimetric mass loss analysis and cross-sectional scanning electron microscopy (SEM), are fundamentally thwarted by extensive host tissue ingrowth, collagen deposition, and neo-vascularization[18]. Beyond the mid-term 16-week post-implantation milestone, the host muscular tissue completely integrates with the dissolving silk filaments, forming an inseparable tissue-mesh complex. This biological overlapping entirely masks the gravimetric signatures of protein resorption and completely obscures explicit fiber architectures, leaving a critical information gap regarding long-term scaffold integrity[19].

To bypass these formidable physical characterization barriers, the field must transition toward intelligent, data-driven computational methodologies[20, 21]. Advanced probabilistic machine learning algorithms, particularly Gaussian Process Regression (GPR), offer a transformative mathematical framework to deconvolve complex polymer degradation pathways[22]. Unlike rigid parametric or purely linear regression models, GPR excels at processing non-linear, high-dimensional, and characteristically small-sample biomedical datasets while uniquely delivering a quantified measure of predictive uncertainty to guard against overfitting[23]. Furthermore, establishing such a framework necessitates successfully bridging the inherent kinetic mismatch that exists between accelerated laboratory testing and complex physiological conditions. By feeding the GPR architecture with high-resolution, normalized in vitro multi-parameter fingerprints—encompassing time-series fiber diameters, gravimetric matrix loss, and the surprisingly conserved molecular structures validated via Fourier-transform infrared spectroscopy (FTIR)— the model can systematically learn the intrinsic material regression rules[24]. By subsequently utilizing in vivo mechanical destructive data as chronological anchor points, the high-resolution in vitro model can be dynamically calibrated onto the timeline of a living organism. This elegant calibration effectively accounts for the complex, bi-phasic in vivo mechanical trajectory, wherein an initial degradation-driven failure phase is sequentially counterbalanced by a secondary mechanical recovery phase driven by biomechanical synergy with the nascent muscular tissue.

In this study, we developed an intelligent, semi-empirical GPR-driven framework designed to systematically decode the “invisible” in vivo degradation trajectory of KSM implants. Importantly, despite the confounding premature fluorescence quenching observed in our platform, this investigation represents the pioneering attempt utilizing optical bioimaging to visually map the complete chronological breakdown of the scaffold’s peripheral boundary layer in vivo, providing indisputable topochemical evidence that outer functionalized envelopes erode rapidly prior to internal silk core dissociation. More crucially, our GPR framework elegantly resolves the perennial technical dilemma of tissue-mesh overlapping. While the absolute physical separation of degraded silk fibers from the surrounding adhered host musculature remains a persistent technical impossibility for traditional gravimetric assays, our computational model successfully addresses this limitation. By mathematically decoupling the intrinsic polymer resorption kinetics from confounding tissue ingrowth signals, the framework achieves the first-of-its-kind lifetime prediction of the bare scaffold’s structural fate in a pristine, non-adhered state, establishing a robust, highly adaptable digital twin platform for the clinical optimization of degradable biomaterials.

## Materials and methods

### 2.1 Materials

#### 2.1.1 Fabrication and Sterilization of Knitted Silk Mesh (KSM)

The Knitted Silk Mesh (KSM) scaffolds are engineered from medical-grade natural silk filaments (22 denier; Beifute, China). Utilizing a computer-controlled Karl Mayer DN21 warp knitting machine, the silk threads are processed under precise tension and winding speeds to ensure structural uniformity. To stabilize the resulting mesh, a thermal annealing process is conducted at 150 °C for 30 minutes in a dry air environment. This specific fabrication protocol is protected under Chinese invention patent ZL 202510135553.6. For downstream biological applications, including cell culture and in vivo implantation, the meshes undergo a rigorous two-step sterilization process: autoclaving (121.3 °C, 103.4 kPa for 30 min) followed by UV irradiation.

#### 2.1.2 Enzyme

The specific type and concentration of the enzyme used for degradation characterization were determined through preliminary screening experiments. All enzymes were purchased from Aladdin. For the formal degradation study, knitted silk meshes (KSM) were incubated at 37°C in a phosphate-buffered saline (PBS) solution containing 10 mg/mL α-chymotrypsin. To maintain constant enzymatic activity, the incubation medium was replenished every three days throughout the designated time intervals (1, 2, 4, 8, 16, and 32 weeks). The extent of degradation was quantified by calculating the percentage of mass loss for each sample group. This specific enzymatic system was selected based on its ability to effectively simulate the proteolysis process within the degradation model.

#### 2.1.3 In vitro degradation and system

Knitted silk meshes (N=3 per group) were incubated at 37°C in a phosphate-buffered saline (PBS) solution containing 10mg/mL α-chymotrypsin. At predetermined time intervals, the specimens were retrieved, rinsed thoroughly with deionized water to terminate enzymatic activity, and subsequently subjected to multimodal characterization: (i) morphological assessment via scanning electron microscopy (SEM); (ii) gravimetric analysis to determine residual mass; (iii) mechanical evaluation using a universal testing machine; and (iv) molecular analysis via Fourier transform infrared spectroscopy (FTIR).

#### 2.1.4 Fluorescent Labeling of Knitted Silk Meshes

To track the degradation process, the knitted silk meshes (KSM) were covalently conjugated with Cyanine7.5 NHS ester (Cy7.5-NHS). Stock solutions of Cy7.5-NHS and 4-dimethylaminopyridine (DMAP) were independently prepared in anhydrous dimethyl sulfoxide (DMSO) at a concentration of 10 mg/mL. A staining working solution was then formulated by diluting appropriate aliquots of the stock solutions into a carbonate buffer (pH=8.5), ensuring the final volumetric concentration of DMSO was maintained below 0.5% to preserve protein conformation.

Subsequently, four KSM specimens were immersed in 10 mL of the prepared staining solution. The reaction was allowed to proceed at 25°C under constant agitation (100 rpm) for 24 hours with strict protection from light. Following the incubation, the meshes underwent a rigorous purification protocol, consisting of sequential washing steps with phosphate-buffered saline (PBS) containing 0.1% Tween-20 (PBST) and pure PBS, respectively, accompanied by centrifugation to eliminate unbound fluorophores and residual reagents. Finally, the labeled meshes were desiccated at room temperature under dark conditions for 24 hours, and subsequently sealed and stored at 4°C in the dark for further characterization.

### 2.2. Methods

#### 2.2.1 Characterization of Scanning Electron Microscopy (SEM)

Scanning electron microscopy (SEM) images were obtained by a field emission scanning electron microscope (Quanta FEG250) at an operating voltage of 3 kV. Before measurements, the degraded KSM were cleaned and dried totally prior to SEM observation. The samples were sputtered with an approximately 10 nm thick Au layer using an EM SCD 500 auto fine coater at a current of 20 mA for 2 min.

#### 2.2.2 Mass Loss Measurement

At each designated time point, the degraded knitted silk meshes (KSM) were retrieved and subjected to a rigorous cleaning protocol to eliminate residual α-chymotrypsin. Specifically, the samples were immersed in a solution of 0.3% detergent and 0.02 M sodium carbonate at 90°C for 30 min. Periodic agitation was performed every 10 min to facilitate detergent suspension and ensure the complete removal of surface-absorbed protein aggregates. Following purification, each mesh was rinsed in Milli-Q water for 90 s under constant agitation. To ensure consistent weight measurement, the samples were blotted on lint-free towels until no visible watermarks remained, followed by thorough dehydration in a convection oven at 60°C for 6 h. The gravimetric mass loss was then calculated for each sample group.

#### 2.2.3 Mechanical Characterization

The mechanical properties of the knitted silk meshes (KSM) were evaluated via uniaxial tensile tests using a universal testing machine AGS-X equipped with a 1000 N load cell. To ensure consistency across all experimental groups, specimens were precisely die-cut into a rectangular geometry with dimensions of 1 × 2 cm^2^.

During the measurement, the samples were secured between the pneumatic grips with an initial gauge length of 10 mm. Tensile loading was applied at a constant crosshead speed of 0.05 mm/s until physical failure occurred. The elongation at break (ε_b_, %), defined as the maximum strain before specimen rupture, was recorded as the primary indicator to characterize the structural ductility and mechanical integrity of the meshes. At least five replicates were tested for each group to ensure statistical significance, and the results were expressed as mean ± SD.

#### 2.2.4 FTIR Characterization

Infrared spectra of KSM were measured by using FTIP spectroscopy (Bruker Tensor 27). All the spectra were obtained at different peratures with 32 scans and a resolution of 2 cm^-1^ in the range of 400 cm^-1^ to 4000 cm^-1^.

#### 2.2.5 Establishment of Partial Abdominal Wall Defect Model and Mesh Implantation

Sprague-Dawley rats were initially anesthetized via inhalation of 5% isoflurane at a flow rate of 5 cc/min and subsequently maintained under a deep anesthetic state with 2% isoflurane at 2 cc/min. Following immobilization in a supine position, the abdominal surgical field was shaved and prepared aseptically with three sequential applications of 75% ethanol.

A 3cm longitudinal incision was made along the lower abdomen, followed by blunt dissection of the skin and subcutaneous tissues to expose the underlying musculature. To establish a partial-thickness abdominal wall defect measuring 1 × 1cm^2^, a designated portion of the external and internal oblique muscles lateral to the lineal alba was surgically excised, while meticulously preserving the underlying transversus abdominis muscle and the peritoneum.

Under strict light-shielding conditions to prevent fluorophore photobleaching, a matching fluorescently labeled knitted silk mesh was adapted to the defect site. The scaffold was secured to the surrounding healthy tissue using 5-0 absorbable sutures via a tension-free interruption technique. The anatomical layers of the abdominal wall and skin were then closed sequentially, followed by topical disinfection. Intraoperative blood loss was minimal, ranging between 1 and 2mL. Postoperatively, analgesia was administered via ibuprofen-infused gel blocks to ensure animal welfare.

#### 2.2.6 In Vivo Fluorescence Tracking via Imaging System

To dynamically monitor the chronological fate of the scaffolds in vivo, non-invasive fluorescence imaging was performed using an IVIS Spectrum in vivo imaging system (PerkinElmer, USA). At predetermined longitudinal time points (weeks 0, 1, 2, 4, 8, 12, and 24 post-surgery), the rats were anesthetized with 2% isoflurane and positioned symmetrically within the imaging chamber.

Based on the spectral properties of the conjugated Cyanine7.5 fluorophore, fluorescence signals were captured using an excitation wavelength of 750 nm and an emission wavelength of 820 nm. Consistent exposure parameters were maintained across all imaging sessions to ensure comparative precision.

Quantitative assessment of the fluorescence attenuation was conducted using Living Image 4.4 software. The background fluorescence from the surrounding native tissue was subtracted to yield the net signal intensity for further degradation kinetic modeling.

#### 2.2.7 Ethical statement and preclinical model for in vivo degradation

All animal experiments were approved by the Institutional Animal Care and Use Committee (IACUC) of Peking University People’s Hospital (Approval No. 2024PHE033). Female Sprague-Dawley rats (7 weeks old, weighing 200 ± 20 g; Charles River Laboratories, China) were acclimatized to specific pathogen-free (SPF) conditions for 7 days prior to surgery. A total of 18 rats were randomly divided into six time-point groups (n = 3 per group). Anesthesia was induced by inhalation of 5% isoflurane/95% O_2_ at a flow rate of 5 L/min and maintained at 2.5%. After shaving and disinfection with iodine, bilateral partial-thickness defects (1 × 2 cm^2^) were created by excising the external and internal oblique muscles 0.5 cm lateral to the linea alba. Mesh implantation was performed using an intra-abdominal overlay technique, and the mesh was secured tension-free to the surrounding fascia with interrupted 5-0 non-absorbable sutures. Postoperatively, the animals were housed individually and routinely observed. The meshes were harvested at 1, 2, 4, 8, 16, and 32 weeks after surgery.

#### 2.2.8 A GPR-Driven Semiempirical Predictive Framework via Time-Scale Calibration

To quantitatively deconvolve and mathematically reconstruct the long-term in vivo degradation kinetics of the KSM, a hybrid data-driven predictive framework was established, shown in Figure 1. This semi-empirical model synergistically integrates multi-parameter in vitro time-series regression with sparse in vivo mechanical anchor points.

**Fig.1.**
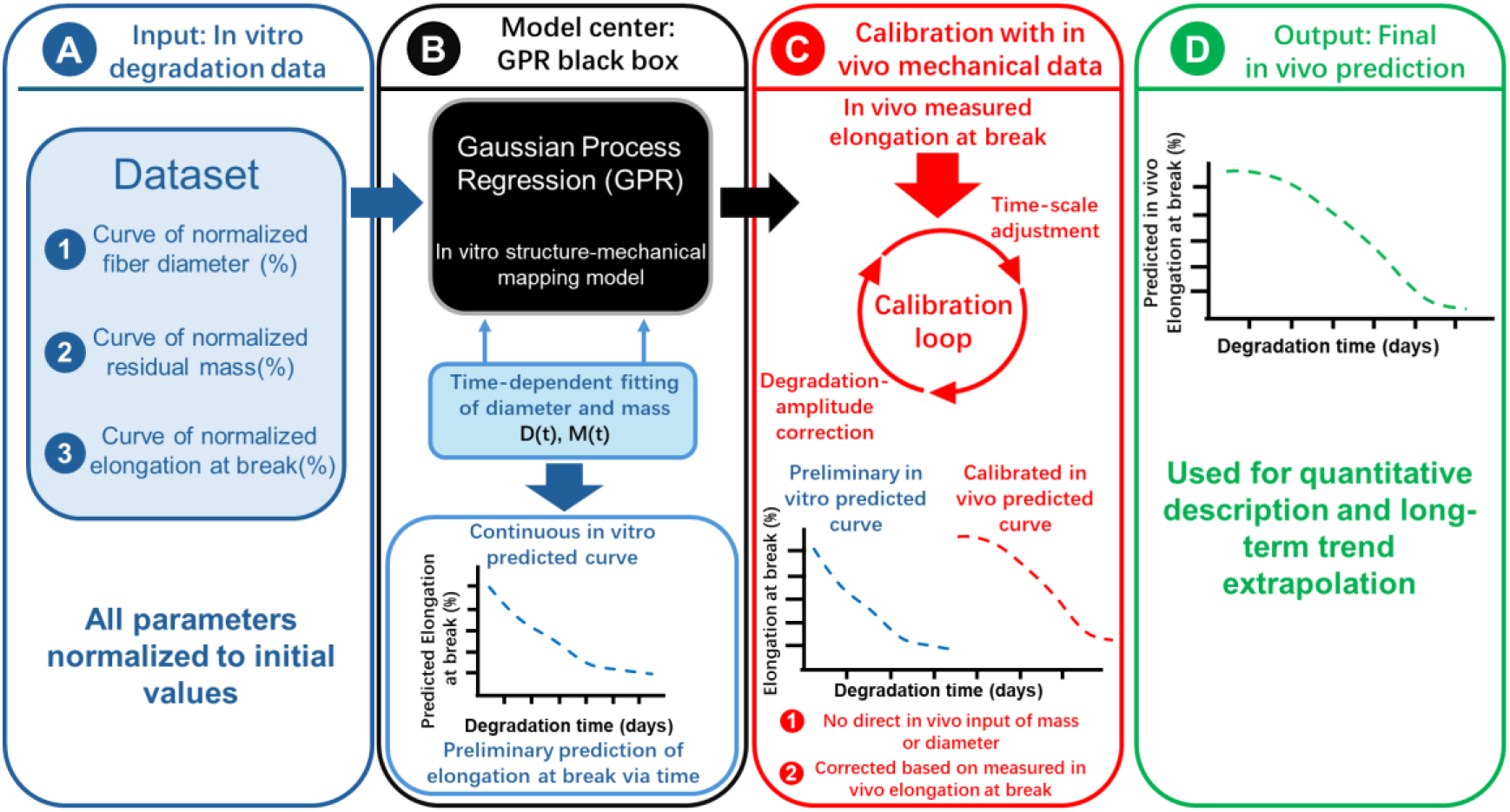
Schematic workflow of the GPR-driven semi-empirical predictive framework. (A) High-dimensional in vitro dataset construction encompassing normalized fiber diameter, residual mass, and elongation at break. (B) Training of the Gaussian Process Regression (GPR) model to establish the structure-mechanical mapping and time-dependent continuous in vitro prediction. (C) Cross-environment calibration loop utilizing sparse in vivo mechanical anchor points via time-scale adjustment and degradation-amplitude correction. (D) Final calibrated in vivo structural fate output for long-term quantitative trend extrapolation. All curves are represented by dashed lines for illustrative purposes only and do not represent actual experimental data.

Initially, high-resolution multi-dimensional degradation descriptors— encompassing effective fiber diameter, cumulative residual mass, and elongation at break—were harvested from the in vitro accelerated assays at discrete longitudinal intervals (weeks 1, 4, 8, 16, and 32). Fig. 1(A) schematically illustrates the parameters comprising the input dataset, while the corresponding experimental curves are provided in Fig. 3. To eliminate inter-sample stochastic variability and initial physical boundaries, all characterization metrics were subjected to feature normalization relative to their pristine, non-degraded states, shown in Fig. 1A.

Utilizing these standardized profiles, a Gaussian Process Regression (GPR) algorithm was deployed to construct a non-linear structural-mechanical mapping model. Within this architectural matrix, the degradation duration (t), normalized fiber diameter (D), and normalized residual mass (M) were designated as high-dimensional input features, while the normalized elongation at break (εb) served as the ultimate target output vector. GPR was specifically selected due to its exceptional capability to handle small-sample biological datasets without sacrificing mathematical rigor, alongside its intrinsic capacity to output quantified predictive confidence intervals (CI).

To resolve the discrete data constraints, the temporal decay pathways of both D and M were mathematically fitted into continuous time-dependent empirical functions and back-propagated into the primary GPR core. This iterative loop yielded a continuous, uninterrupted in vitro structural-mechanical deterioration curve.

## 3. Results and Discussion

### 3.1 In Vivo Spatiotemporal Fluorescence Decay Kinetics and Structural Disintegration Hierarchy

The real-time in vivo degradation kinetics of the KSM scaffolds were longitudinally monitored via non-invasive fluorescence imaging over a 32-week tracking matrix (Fig. 2A), with the corresponding total fluorescence intensity quantified in Fig. 2B. Chronologically, the fluorescence signal exhibited an anomalous yet distinct bi-phasic trajectory. During the early post-implantation phase (weeks 1 to 4), a transient escalation in fluorescence intensity was recorded. This initial signal spike can be synergistically rationalized by the spatial relaxation of the fluorophores. When the pristine KSM scaffolds were statically incubated in the cyanine dye solution during the initial labeling process, an ultra-high local concentration of dye molecules accumulated within the localized domains. This close molecular proximity induced a severe fluorescence self-quenching state. Upon implantation, host fluid infiltration and initial matrix swelling expanded the inter-fluorophore spatial distance, thereby relieving the self-quenching constraints (de-quenching) and precipitating the observed early-stage fluorescence surge.

**Fig. 2.**
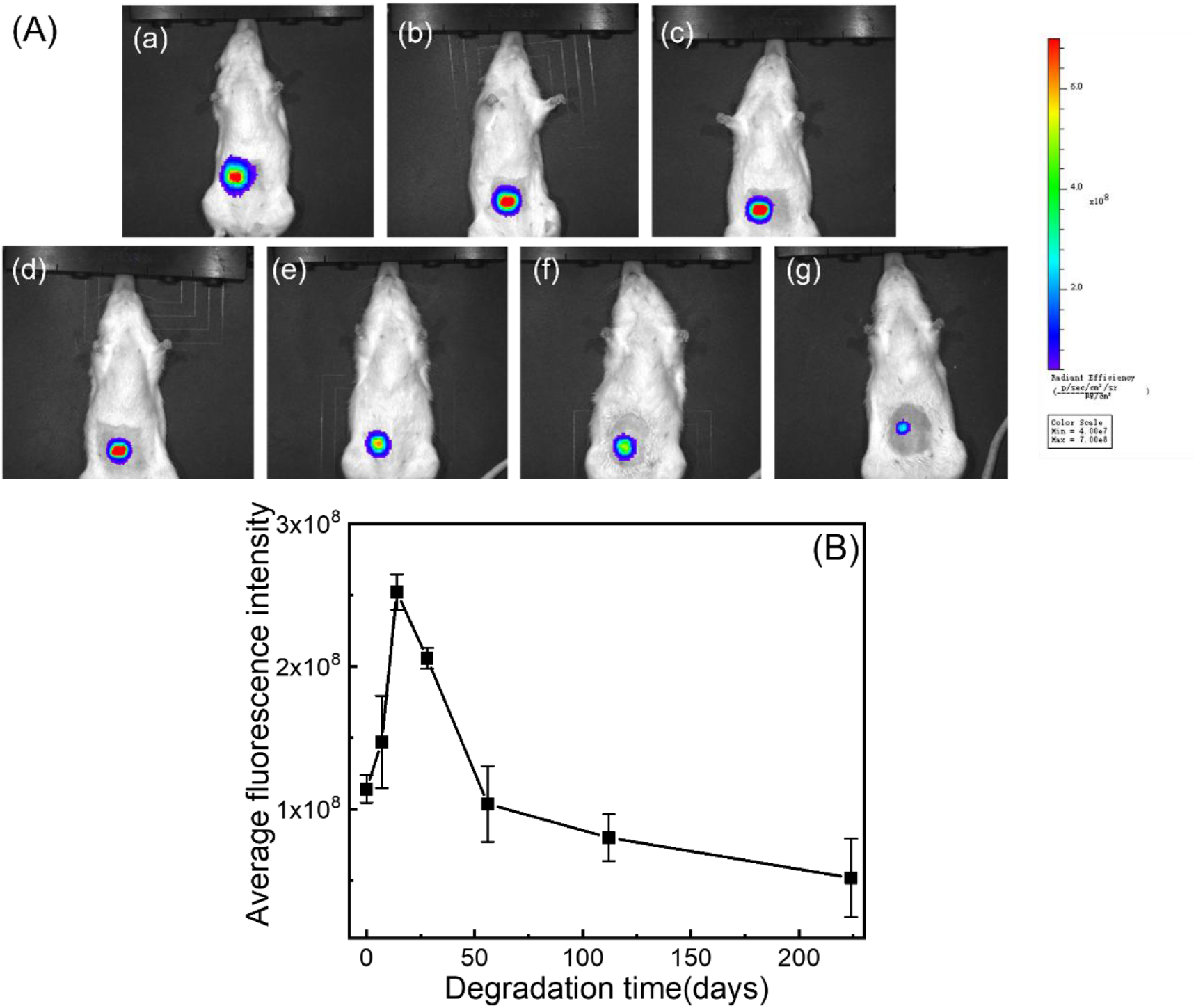
In vivo fluorescence tracking and quantitative fluorescence intensity of knitted silk meshes (KSM) in a rat model. (A) Representative in vivo fluorescence images tracking the degree of degrading at 7 sequential post-implantation checkpoints (0, 1, 2, 4, 8, 16 and 32 weeks). (B) Quantitative line graph of the corresponding average fluorescence intensity over the 32-week degradation period.

As degradation progressed into the mid-term phase (weeks 8 to 16), a precipitous decline in fluorescence intensity was captured, which aligned with the macroscopic mass loss trends and mechanical regression curves. By the late-stage checkpoints (weeks 24 to 32), the remaining fluorescence signal rapidly plateaued and eventually diminished to the baseline threshold, rendering the scaffolds virtually “invisible” under optical bioimaging. Paradoxically, parallel gravimetric analyses and macroscopic tissue examinations corroborated that a substantial portion of the KSM structural framework still retained its physical integrity at these advanced time points.

This critical kinetic mismatch reveals a profound structural degradation hierarchy rather than an experimental artifact. Due to the high steric hindrance and dense packing architecture of the pristine fibroin bundles, the initial dye conjugation was restricted to a “peripheral labeling” mode, wherein the fluorophores were predominantly grafted onto the outer functionalized shell, leaving the internal core matrix un-dyed. Consequently, while the premature loss of fluorescence signals might superficially mimic complete material dissolution, it paradoxically provides definitive, exclusive evidence for the complete topochemical erosion of the scaffold’s peripheral boundary layer. Taken together, these findings elucidate a sequential in vivo degradation mechanism of KSM: a rapid, enzyme-driven cleavage initiated at the outer functionalized envelope, which sequentially precedes the subsequent structural dissociation, unraveling, and eventual breakdown of the internal silk fibroin cores.

### 3.2. Multi-Parameter Evaluation of In Vitro Degradation: Mass Loss, Diameter Reduction, and Mechanical Decay

To quantitatively map the structural regression of the scaffold, the chronological kinetics of cumulative mass loss and effective fiber diameter were systematically evaluated, seen in Fig. 3. It must be noted that due to the extensive tissue encapsulation and inseparable host-material integration observed in vivo after week 16, shown in Fig. 5, precise isolation of the remnant meshes without sacrificing structural fidelity was unfeasible. Consequently, the longitudinal gravimetric profiles were successfully established utilizing the in vitro accelerated degradation model as a high-resolution proxy to delineate the intrinsic resorption velocity of the silk framework.

**Fig. 3.**
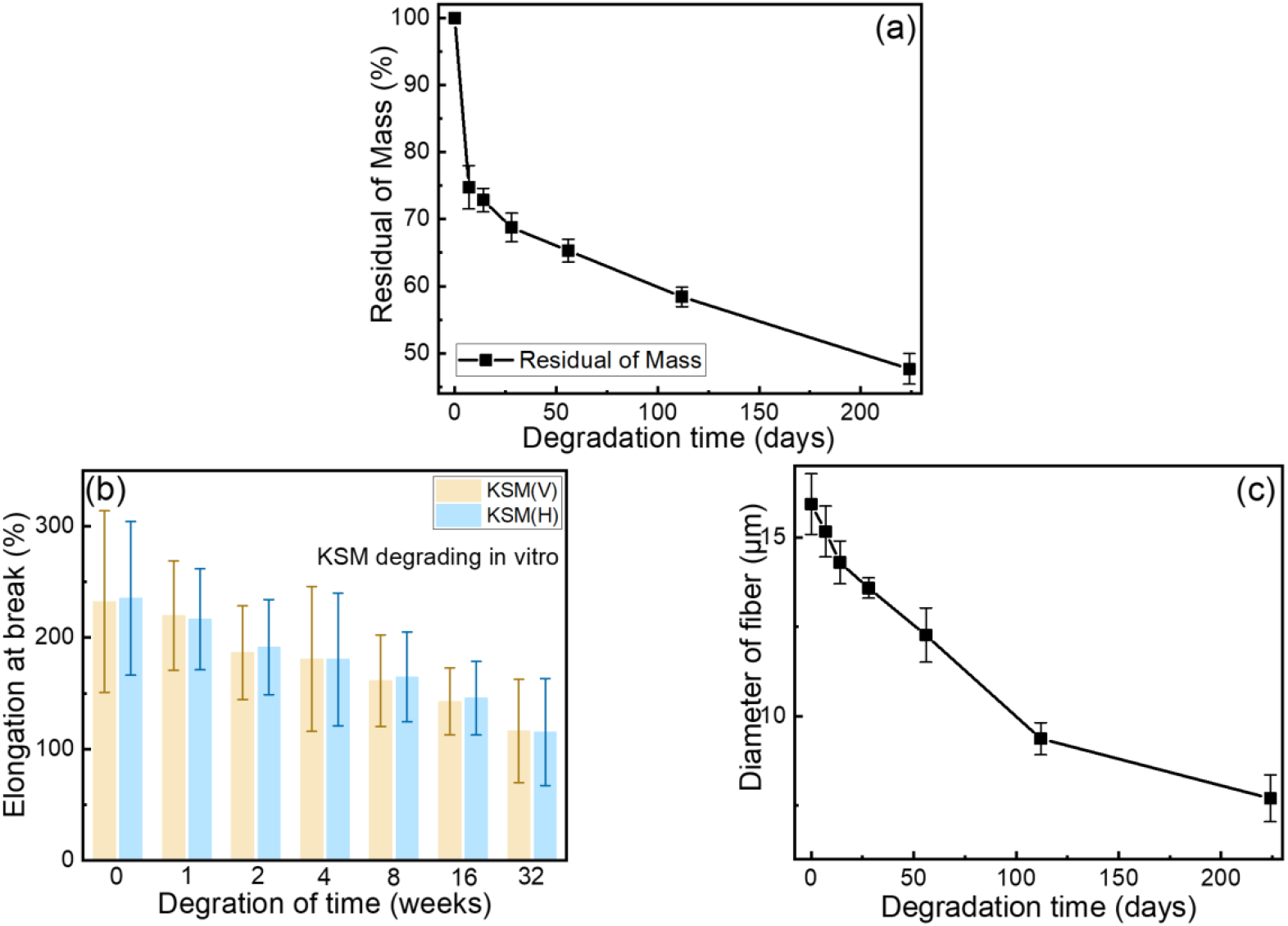
In vitro enzymatic degradation kinetics of knitted silk meshes (KSM). (a) Chronological profiling of the normalized residual mass percentage over the incubation period. (b) Elongation at break along both longitudinal and transverse directions. (c) Microstructural evolution tracking silk fibroin fiber diameters quantified via scanning electron microscopy (SEM) analysis.

The chronological mass loss profile unfolded via a distinct bi-phasic tracking behavior, characterizing a classic topochemical erosion mechanism. In the opening phase of enzymatic incubation (weeks 1–8), the scaffold underwent a rapid, highly accelerated mass attrition. This initial gravimetric drop correlates dynamically with early-stage SEM evidence, which revealed that the hierarchical disintegration of macro-scale multi-fiber bundles into individualized, free-standing single strands drastically maximized the effective surface area exposed to enzymatic attack, prompting aggressive hydrolytic cleavage of the highly accessible, disordered amorphous domains. Intriguingly, past the 8-week threshold, the kinetics of mass resorption transitioned from a sharp decline into a dampened, near-linear decay. This late-stage deceleration stabilized into a steady downward trajectory, signaling that the enzymatic attack had shifted from the easily digestible outer envelope to the highly resilient, densely packed crystalline β-sheet cores.

This quantitative kinetic shift provides profound insights when cross-examined with the prior morphological transformations. The proliferation of localized surface pits and craters did not immediately trigger a massive bulk weight loss. More importantly, the large-scale lamellar delamination and membrane peeling observed in the advanced stages did not precipitate a catastrophic acceleration in mass attrition, but rather manifested primarily as a severe topological transformation. This apparent scaling mismatch can be comprehensively explained by the physical state of the exfoliated fragments. Concurrently, the temporal evolution of the effective fiber diameter displayed a highly synchronized downward trend with the initial weight decay, seen in Fig. 3c. The continuous thinning of the fiber cores liberated finer sub-fibrils, leading to early mass shedding. However, during the advanced stages (beyond 16 weeks), although large sections of the proteinaceous sheets underwent physical delamination and separated from the parental fiber core, these detached structural components remained spatially confined and weakly entrapped within the intricate, interlocking knitted topology of the mesh. Because these sloughed sheets were not fully liberated or dissolved into the surrounding bulk medium, they still registered gravimetrically during sampling. This spatial entrapment artifact induced a subtle, characteristic deviation in the late-stage mass loss curve, causing it to decouple slightly from the severe physical disintegration observed under high-vacuum SEM.

### 3.3. Morphological Evolution and Microstructural Disintegration of KSM

The progressive in vitro degradation trajectory of the knitted silk meshes (KSM) under a constant enzymatic concentration was chronologically elucidated via SEM analysis, shown in Fig. 4. As illustrated in the micrographs, the morphological regression of the scaffold followed a multi-stage, topochemical erosion pathway.

**Fig. 4.**
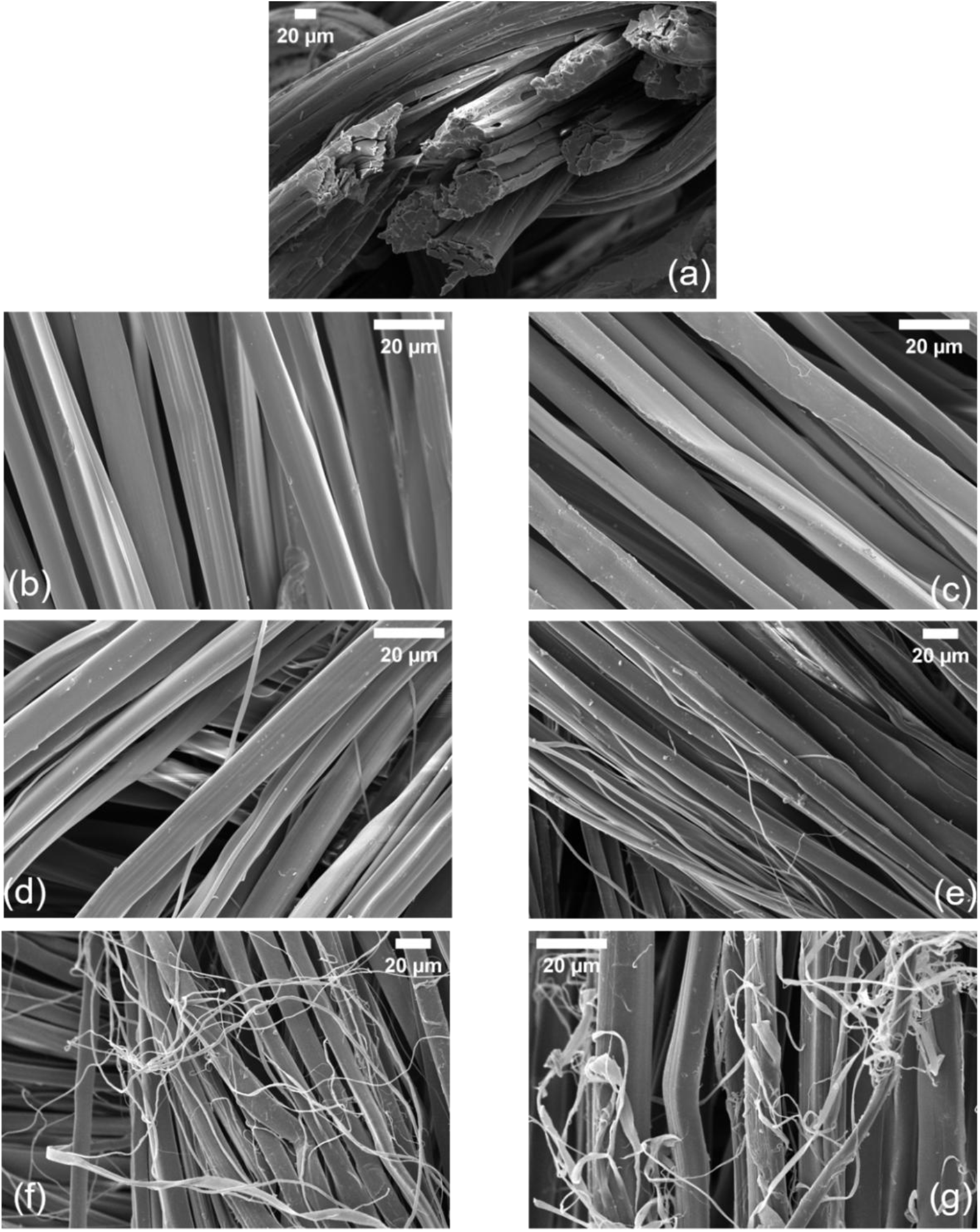
SEM micrographs showing the morphological evolution of knitted silk meshes (KSM) during 32 weeks of in vitro enzymatic degradation. (a) to (g) chronologically represent the sequential structural changes at 0, 1, 2, 4, 8, 16 and 32 weeks of incubation, illustrating the transition from fiber bundle dissociation to lamellar delamination.

In the initial phase of incubation, seen in Fig. 4b-c, the degradation initiated with the structural dissociation of the highly ordered multi-fiber bundles. The constituent silk strands gradually untangled and isolated into discrete, individualized micro-fibers. Distinctly, despite this individualization of the strands, the macro-scale mesh retained its integral, porous net-like framework rather than suffering a catastrophic collapse. This structural stability is primarily attributed to the interlocking knitted architecture and local knotted junctions, which effectively confined the separated strands and maintained the spatial topology of the scaffold during early-stage mass loss.

As the exposure time to α-chymotrypsin was prolonged, shown in Fig. 4d-e, the erosion extended from the inter-fiber interfaces deeper into the intra-fiber matrices. The surface of the isolated single fibers began to undergo pronounced micro-fibrillation, characterized by the physical detachment of finer, sub-micron protein fibrils from the parental fiber trunks. Concurrently, localized morphological defects, including distinct surface pits, craters, and structural indentations, began to proliferate along the longitudinal axes of the fibers.

With a further increase in degradation duration shown in Fig. 4f-g, this micro-fibrillation amplified drastically. A high density of sub-fibrils exfoliated from the matrix, accompanied by a noticeable and continuous reduction in the effective fiber diameter. Most notably, in the advanced stages of degradation (beyond 16 weeks, Fig. 4g), the erosion mode transitioned from localized pitting to severe lamellar delamination. Entire polymeric sheets and bulk proteinaceous fragments were observed to strip and peel away from the residual fiber cores, akin to the shedding of a structural membrane. This catastrophic physical disintegration created a interconnected network of micro-voids, which not only directly accounted for the macro-scale mass attrition but also acted as pervasive stress concentrators, explaining the subsequent decay in tensile properties.

### 3.4. Chronological In Vivo Morphological Regression and Tissue Integration Interplay

To further deconstruct the physiological fate of the scaffold, the chronological morphological alterations of the KSM retrieved from the murine model were systematically investigated (Figure 2). Intriguingly, the in vivo degradation modality exhibited a distinct divergence from the in vitro fibrillation-dominated pathway.

In the early post-implantation phase (Figure 5a-b), the KSM specimens showed scarcely any detectable micro-fibrillation or individualization of single silk strands. Instead, the morphological topography was characterized by a high density of localized, severe perforations, deep pits, and micro-cavities along the fiber boundaries. This unique phenomenon suggests that in vivo degradation is primarily governed by localized cell-mediated enzymatic erosion, where host inflammatory cells (e.g., macrophages) adhere to the material surface and secrete concentrated proteolytic enzymes, generating discrete pocket-like defects rather than uniform stripping.

**Fig. 5.**
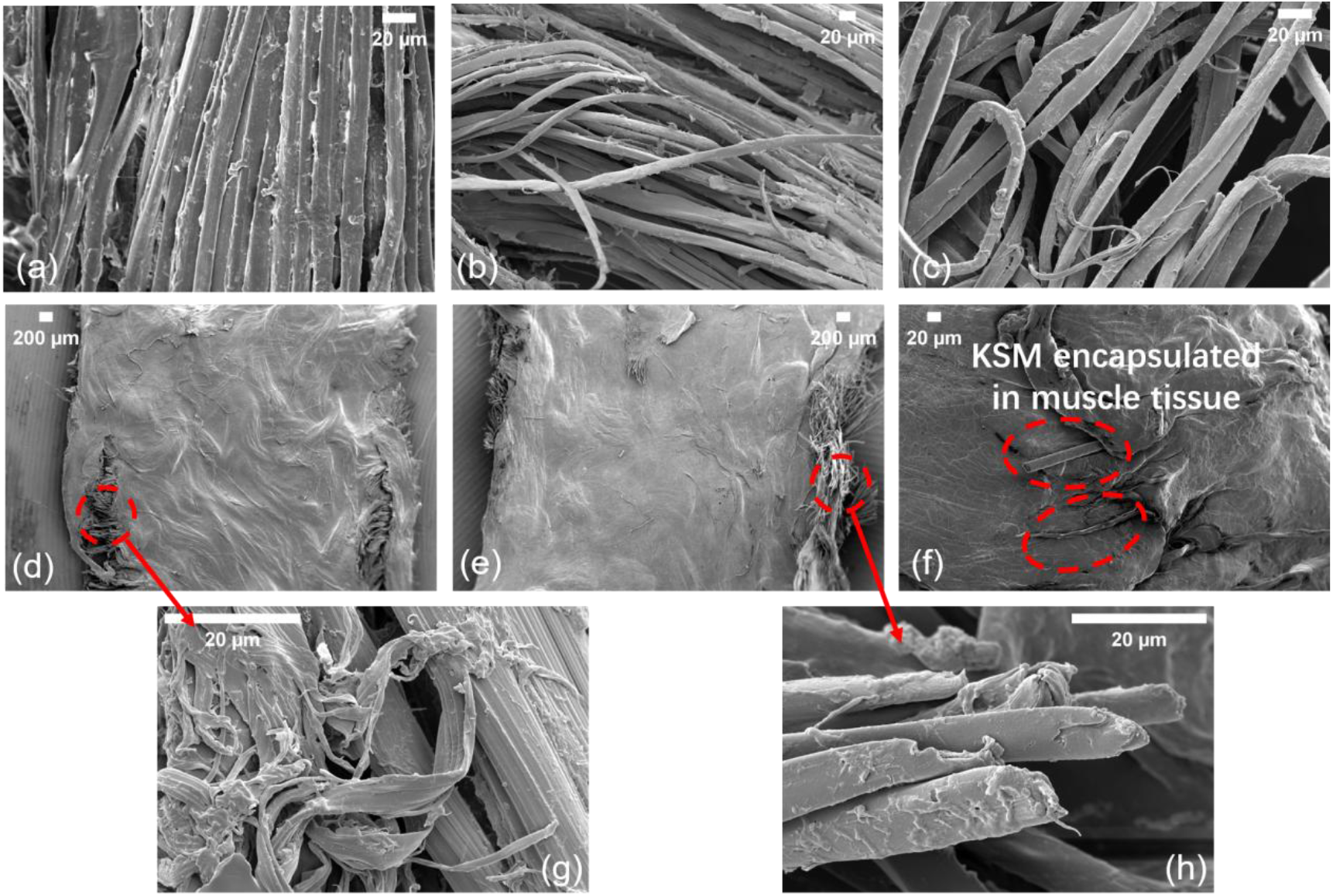
SEM micrographs tracking the in vivo degradation and tissue integration of knitted silk meshes (KSM) in a rat model. (a) to (f) chronologically display the explant morphology at 0, 1, 4, 8, 16, 24, and 32 weeks post-implantation, demonstrating the cellular erosion process and eventual integration with the host muscular tissue. The dashed circles provide magnified views highlighting specific microstructural alterations: (g) demonstrating macro-scale extensive lamellar delamination, and (h) exhibiting severe superficial topochemical erosion.

As the implantation duration extended into the mid-term phase (Figure 5b), the progressive accumulation and coalescence of these localized defects induced a structural transition. The micrographs captured large-scale lamellar delamination and massive bulk matrix collapse. Wide areas of the proteinaceous sheets appeared to disintegrate and slough off concurrently, creating interconnected macro-porous networks and structural voids. This aggressive structural regression provided definitive morphological evidence that the resorption of the silk backbone was actively and continuously proceeding in vivo.

Most notably, in the long-term degradation phase (beyond 16 weeks, Fig 5c), the dynamic interplay between material resorption and host remodeling reached an advanced stage. The KSM became meticulously encapsulated and intertwined with the nascent rat pelvic floor musculature. Because of this extensive cell infiltration and neo-tissue deposition, the scaffolds formed an inseparable dense tissue-mesh complex that could not be isolated without causing irreversible tearing to the residual matrix. Consequently, the resulting SEM micrographs appeared characteristically obscured and masked by the overlying host extracellular matrix. Nevertheless, through sparse windows of exposed structural components (Fig. 5d), it was highly evident that the majority of the remaining silk backbones had anatomically integrated and fused with the surrounding muscular tissues. The boundaries between the synthetic template and native host matrix became virtually indistinguishable, fundamentally validating the exceptional histocompatibility and tissue-inductive properties of the KSM.

### 3.5. Mechanistic Divergence Governing In Vitro and In Vivo Degradation Kinetics

A comparative cross-examination of the in vitro and in vivo SEM profiles reveals that the degradation velocity of the KSM in vivo significantly outpaces that observed under controlled in vitro conditions. This pronounced kinetic acceleration and the divergent degradation modalities can be synergistically rationalized by the highly dynamic physiological environment and the complex material-host tissue interplay.

First, the metabolic homeostasis of the host environment plays a decisive role. In the in vitro system, despite the periodic replacement of α-chymotrypsin, the enzymatic activity inevitably experiences a temporal decay, and the static incubation system suffers from localized accumulation of acidic degradation byproducts that can inhibit sustained enzyme efficiency. Conversely, the in vivo microenvironment within the rat model undergoes continuous metabolic clearance and physiological circulation. This process ensures a perpetual supply of active endogenous proteolytic enzymes (such as matrix metalloproteinases) and a highly stabilized physiological $pH$, maintaining a constant and aggressive degradation pressure on the implant.

Second, the spatial boundary conditions of the degradation interface differ fundamentally. In the in vitro framework, the KSM is entirely submerged in the enzymatic medium, resulting in isotropic erosion that operates predominantly at the scale of individualized single silk fibers (characterized by uniform micro-fibrillation). In sharp contrast, in vivo degradation progresses synchronously with host tissue ingrowth and neo-muscularization. Owing to the exceptional biocompatibility of the silk fibroin matrix, the newly regenerated muscular tissue adheres tightly to and wraps around the silk strands. This bio-interface seals off portions of the fibers, creating localized micro-domains where host inflammatory cells (e.g., macrophages) adhere and release concentrated, high-potency enzymes. This contact-dependent, cell-mediated attack explains why the in vivo morphology is dominated by deep pits and macroscopic lamellar shedding rather than uniform fiber thinning.

Finally, the influence of host physiological locomotion cannot be overlooked. Unlike the static or low-shear immersion in vitro, the implanted KSM in vivo is subjected to continuous cyclic mechanical loading generated by the rat’s respiration, ambulation, and intra-abdominal pressure. This sustained in-service mechanical stress induces micro-deformation within the knitting knots and accelerates the propagation of micro-cracks and structural defects created by enzymatic erosion. The coupling effect of mechanical fatigue and active biochemical cleavage dynamically accelerates the macro-scale structural breakdown of the mesh in vivo.

These profound discrepancies thoroughly invalidate the feasibility of utilizing simple linear scaling to extrapolate in vivo fate from in vitro benchmarks. This, in turn, provides a compelling justification for our deployment of the Gaussian Process Regression (GPR) framework, which leverages sparse in vivo mechanical anchor points to dynamically calibrate the non-linear, multi-scale degradation trajectory of the biomaterial.

### 3.6. FTIR Spectroscopic Analysis of Chemical Structural Evolution During Degradation

The chronological FTIR spectra revealed a remarkably resilient chemical profile across all predetermined incubation intervals up to 32 weeks, shown in Fig. 6. Specifically, the characteristic absorption fingerprints of silk fibroin—including the Amide I band (1620–1650 cm^-1^) governed by C=O stretching, the Amide II band (1515–1530 cm^-1^) associated with N-H bending, and the Amide III band (1230–1260 cm^-1^)—exhibited no noticeable chronological shifts, peak broadening, or the emergence of novel chemical functionalities.

**Fig. 6.**
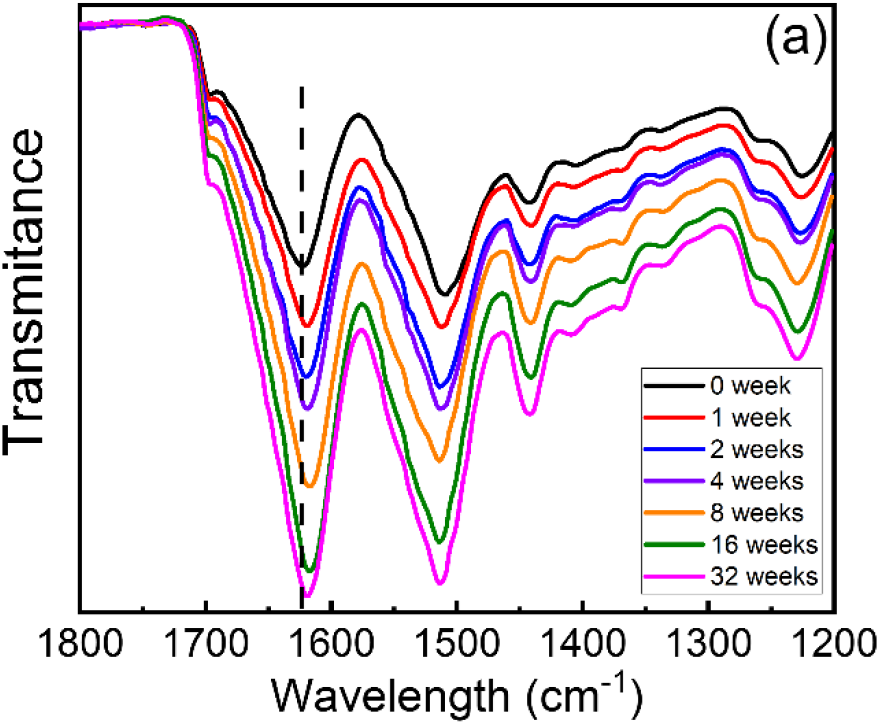
FTIR spectra of knitted silk mesh incubated in 10mg/mL α-chymotrypsin for 0, 1, 2, 4, 8, 16 and 32 weeks.

This spectral invariance firmly demonstrates that even under sustained, aggressive proteolytic attack by α-chymotrypsin, the minimum structural units and primary chemical essence of the residual silk matrix remain highly conserved and completely unaffected by macro-scale physical disintegration.

### 3.7. GPR-Based Prediction of In Vivo KSM Degradation

A critical bottleneck in evaluating the in vivo degradation kinetics of tissue-engineered scaffolds lies in the physical confounding caused by rapid host tissue integration. As demonstrated from Week 8 onward, the progressive ingrowth of neo-muscular tissue rendered the complete separation of KSM matrices from the host anatomical site increasingly unfeasible. By inputting the comprehensive in vitro multi-parameter dataset (mass loss, fiber diameter, and elongation at break) and deploying sparse in vivo mechanical anchor points as calibration constraints, the GPR model successfully mathematically decoupled polymer erosion from confounding tissue signatures. Ultimately, the GPR framework successfully isolated the true *in vivo* elongation at break of the bare scaffold by mathematically eliminating confounding host tissue signatures, shown in Fig. 7.

**Fig.7.**
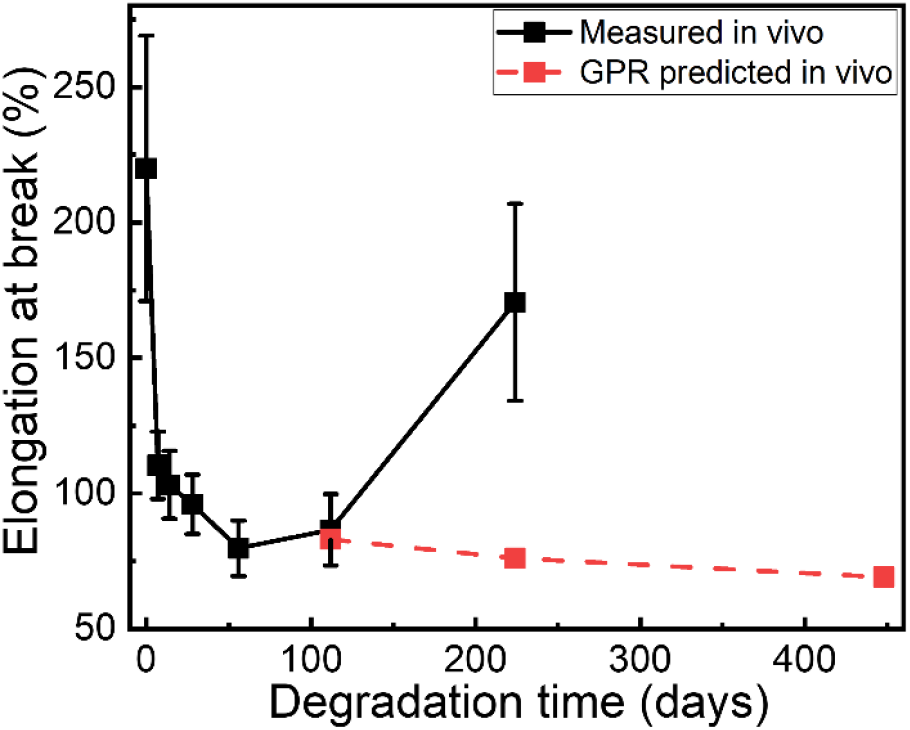
Verification of the GPR model through alignment with experimental mechanical regression. Chronological comparison between the empirically measured in vivo elongation at break (solid line with symbols) and the corresponding Gaussian Process Regression (GPR) fitted trajectory (dashed line). Both datasets are normalized to their initial post-implantation states to eliminate stochastic biological variability, demonstrating the high-fidelity predictive capability of the probabilistic framework in capturing non-linear mechanical decay.

This algorithmic correction effectively recalibrated the in vitro degradation trajectory to fit the in vivo environment, yielding a high-fidelity, unobstructed projection of the standalone scaffold’s long-term in vivo degradation degree. This digital twin methodology offers a powerful and non-destructive paradigm for the lifetime prediction of degradable biomaterials where traditional destructive tracking methods encounter severe boundaries.

## 4. Conclusion

The structural fate of the KSM scaffolds post-implantation presents a complex, multi-layered chronological behavior. Intriguingly, while the baseline fluorescence signal faded prematurely around week 16, this rapid decay should not be dismissed as a mere tracking artifact; rather, it provides exclusive topochemical evidence of the complete erosion of the scaffold’s functionalized outer shell, preceding the internal core dissolution. More crucially, our GPR framework elegantly resolved the perennial dilemma of tissue-mesh overlapping. In empirical settings, completely stripping the degraded silk fibers from the adhered muscular matrix is clinically and technically unachievable. By training the GPR center on multi-modal in vitro fingerprints and incorporating in vivo anchoring trends, the intelligent model managed to ‘virtually dissect’ the tissue-mesh complex. It successfully simulated the degradation curve of the bare mesh template in an ideal, tissue-free microenvironment, bridging the critical characterization gaps and offering a robust digital twin for degradable biomaterials.

## Declarations of Ethics Approval and Consent to Participate

The study protocol has been approved by the Ethical Committee of the Peking University People’s Hospital with an approval number (2024PHE033) and performed according to the Declaration of Helsinki. Ethical Committee of the Peking University People’s Hospital waived need for written informed consents due to the retrospective nature of this study.

## Acknowledgement

This work was financially supported by the National Key R&D Program of China (grant no. 2023YFC2411203), the National Natural Science Foundation of China (grant no. W2511011 and grant no. 22273114), the Strategic Priority Research Program of the Chinese Academy of Sciences (grant no. XDB0770101).

## References

1. Olsen, A.L., et al., Epidemiology of surgically managed pelvic organ prolapse and urinary incontinence. OBSTETRICS AND GYNECOLOGY, 1997. 89(4): p. 501–506.

2. Barber, M.D., Pelvic organ prolapse. BMJ, 2016. 354: p. i3853.

3. Dällenbach, P., To mesh or not to mesh: a review of pelvic organ reconstructive surgery. International Journal of Women’s Health, 2015. 7(null): p. 331–343.

4. Barber, M.D., Mesh use in surgery for pelvic organ prolapse. BMJ, 2015. 350: p. h2910.

5. Brincat, C.A., Pelvic Organ Prolapse: Reconsidering Treatment, Innovation, and Failure. JAMA, 2019. 322(11): p. 1047–1048.

6. Aikman, E.L., L.E. Eccles, and W.L. Stoppel, Native Silk Fibers: Protein Sequence and Structure Influences on Thermal and Mechanical Properties. Biomacromolecules, 2025. 26(4): p. 2043–2059.

7. Hu, D., et al., Silk sericin as building blocks of bioactive materials for advanced therapeutics. Journal of Controlled Release, 2023. 353: p. 303–316.

8. Sahoo, J.K., et al., Silk chemistry and biomedical material designs. Nature Reviews Chemistry, 2023. 7(5): p. 302–318.

9. Yucel, T., M.L. Lovett, and D.L. Kaplan, Silk-based biomaterials for sustained drug delivery. Journal of Controlled Release, 2014. 190: p. 381–397.

10. Altman, G.H., et al., Silk-based biomaterials. BIOMATERIALS, 2003. 24(3): p. 401–416.

11. Xue, Y.C., et al., Interplay of the structures and viscoelastic properties of polyampholyte gels with the interlude of neutral blocks. Soft Matter, 2025. 21(36): p. 7042–7053.

12. Wang, G.K., Y.M. Yang, and D. Jia, Programming viscoelastic properties in a complexation gel composite by utilizing entropy-driven topologically frustrated dynamical state. Nature Communications, 2024. 15(1): p. 3569.

13. Raia, N.R., et al., Characterization of silk-hyaluronic acid composite hydrogels towards vitreous humor substitutes. Biomaterials, 2020. 233: p. 119729.

14. Zhao, Y.H. and D. Jia, Dynamically Hydrogen-Bonded Microphase Separation Enabling Phase Transition in the Gel Composites With Tunable UCST. Advanced Materials, 2026. n/a(n/a): p. e73360.

15. Hong, G., A.L. Antaris, and H. Dai, Near-infrared fluorophores for biomedical imaging. Nature Biomedical Engineering, 2017. 1(1): p. 0010.

16. Frangioni, J.V., In vivo near-infrared fluorescence imaging. CURRENT OPINION IN CHEMICAL BIOLOGY, 2003. 7(5): p. 626–634.

17. Li, M., M. Ogiso, and N. Minoura, Enzymatic degradation behavior of porous silk fibroin sheets. Biomaterials, 2003. 24(2): p. 357–365.

18. Badylak, S.F., D.O. Freytes, and T.W. Gilbert, Extracellular matrix as a biological scaffold material: Structure and function. Acta Biomaterialia, 2009. 5(1): p. 1–13.

19. Pierce, L.M., et al., Biomechanical properties of synthetic and biologic graft materials following long-term implantation in the rabbit abdomen and vagina. American Journal of Obstetrics and Gynecology, 2009. 200(5): p. 549.e1-549.e8.

20. Butler, K.T., et al., Machine learning for molecular and materials science. Nature, 2018. 559(7715): p. 547–555.

21. Suwardi, A., et al., Machine Learning-Driven Biomaterials Evolution. Advanced Materials, 2022. 34(1): p. 2102703.

22. Deringer, V.L., et al., Gaussian Process Regression for Materials and Molecules. Chemical Reviews, 2021. 121(16): p. 10073–10141.

23. Rasmussen, C.E., Gaussian Processes in Machine Learning, in Advanced Lectures on Machine Learning: ML Summer Schools 2003, Canberra, Australia, February 2 - 14, 2003, Tübingen, Germany, August 4 - 16, 2003, Revised Lectures, O. Bousquet, U. von Luxburg, and G. Rätsch, Editors. 2004, Springer Berlin Heidelberg: Berlin, Heidelberg. p. 63–71.

24. Jumper, J., et al., Highly accurate protein structure prediction with AlphaFold. Nature, 2021. 596(7873): p. 583–589.

